# A geometric basis for surface habitat complexity and biodiversity

**DOI:** 10.1101/2020.02.03.929521

**Authors:** Damaris Torres-Pulliza, Maria A. Dornelas, Oscar Pizarro, Michael Bewley, Shane A. Blowes, Nader Boutros, Viviana Brambilla, Tory J. Chase, Grace Frank, Ariell Friedman, Mia O. Hoogenboom, Stefan Williams, Kyle J. A. Zawada, Joshua S. Madin

**Affiliations:** Hawai’i Institute of Marine Biology, University of Hawai’i, Kaneohe, HI, United States; Department of Biological Sciences, Macquarie University, Sydney, NSW, Australia; Centre for Biological Diversity, Scottish Oceans Institute, University of St Andrews, St Andrews KY16 9TH, UK; Australian Centre for Field Robotics, University of Sydney, Sydney, NSW, Australia; German Centre for Integrative Biodiversity Research (iDiv) Halle-Jena-Leipzig, Deutscher Platz 5e, Leipzig 04103, Germany; Department of Computer Science, Martin Luther University Halle-Wittenberg, Am Kirchtor 1, Halle (Salle) 06108, Germany; ARC Centre of Excellence for Coral Reef Studies and College of Science and Engineering, James Cook University, Townsville, Queensland 4811, Australia; Greybits Engineering, Sydney, NSW, Australia

## Abstract

Structurally complex habitats tend to contain more species and higher total abundances than simple habitats. This ecological paradigm is grounded in first principles: species richness scales with area, and surface area and niche density increase with three-dimensional complexity. Here we present a geometric basis for surface habitats that unifies ecosystems and spatial scales. The theory is framed by fundamental geometric constraints among three structure descriptors—surface height, rugosity and fractal dimension—and explains 98% of surface variation in a structurally complex test system: coral reefs. We then show how coral biodiversity metrics (species richness, total abundance and probability of interspecific encounter) vary over the theoretical structure descriptor plane, demonstrating the value of the theory for predicting the consequences of natural and human modifications of surface structure.

## Main text

Most habitats on the planet are surface habitats—from the abyssal trenches to the tops of mountains, from coral reefs to the tundra. These habitats exhibit a broad range of structural complexities, from relatively simple, planar surfaces to highly complex three-dimensional structures. Currently, human and natural disturbances are changing the complexity of habitats faster than at any time in history^1–4^. Therefore, understanding and predicting the effects of habitat complexity changes on biodiversity is of paramount importance^5^. However, empirical relationships between commonly-used descriptors of structural complexity and biodiversity are variable, often weak or contrary to expectation^6–10^. Moreover, there are no standards for quantifying structural complexity, precluding general patterns in the relationship between structure and diversity from being identified in different habitats. We therefore propose a new geometric basis for surface habitats that integrates and standardises existing surface descriptors^8,10^.

In theory, species richness scales with surface area according to a power law^11^. Island biogeography theory articulates that this relationship arises out of extinction and colonization, as larger areas provide larger targets for species to colonize and a greater variety of habitats allowing species to coexist^12^. Our geometric theory builds on these ideas by exploring the notion that habitat surfaces with the same rugosity (defined here as surface area per planar area) can exhibit a range of different forms (Fig. 1). Total surface area is the integration of component areas at the smallest scale (i.e., resolution), but it does not explain how these component areas fold and fill the three-dimensional spaces they occupy. Rather, fractal dimension quantifies space-filling at different scales^13^. Space-filling promotes species co-existence by dividing surface area into a greater variety of structural elements^14^, microhabitats and niches^15^ (e.g., high and low irradiance; small and large spaces; fast and slow flow). This variety of niches allows species to coexist (e.g. different competitors, or predator and prey^16^) and therefore enhances biodiversity^17,18^. We posit that there is a fundamental geometric constraint between surface rugosity and fractal dimension: for a given surface rugosity, an increase in fractal dimension will result in a reduction of the surface’s mean height (Fig. 1). As the basis for a geometric theory, we mathematically derived the trade-off between surface rugosity (*R*), fractal dimension (*D*) and surface height range (Δ*H*) as (see Methods for derivation):

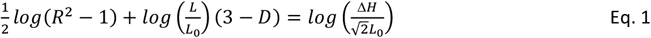

**Fig. 1.**
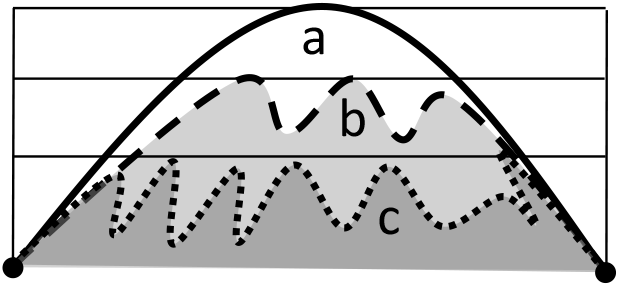
Increasing fractal dimension (i.e., space filling) while keeping surface rugosity constant results in a decline in a surface’s mean height range. A two-dimensional representation of three hypothetical surface habitats with the same surface rugosities (**a**, **b** and **c**). That is, the lengths of the lines **a**, **b** and **c** are the same and occur over the same planar extent (black points). However, line **a** fills less of its two-dimensional space (black rectangle) than does line **c**, and therefore has a lower fractal dimension.

Where *L* is the surface linear extent and *L*_0_ is the resolution (i.e., the smallest scale of observation). *R* and *D* are both dimensionless, with *R* ≥ 1 and 2 ≤ *D* ≤ 3; Δ*H* is dimensionless when standardised by resolution *L*_0_, with 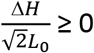. When rugosity is expressed as *R*^2^-1 (with *R*^2^-1 ≥ 0) and height range as 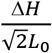, Eq. 1 is a plane equation. Moreover, it is clear that any one of the surface descriptors can easily be expressed in terms of the other two, highlighting that any of the three variables is required, but not sufficient alone, to describe the structural complexity of a surface habitat.

## Results

To test the theory, we examined associations among surface rugosity, fractal dimension and height range across coral reef habitat patches. Coral reefs are ideal ecosystems for testing a theory of surface habitats, because they are structurally complex surface habitats constructed in large part by the reef-building scleractinian corals that, in turn, live upon the habitat (i.e., corals are autogenic ecosystem engineers^19^). Structural complexity affects biodiversity in general^20^ and of coral reefs in particular^21^. Using Structure from Motion (SfM), we estimated surface rugosity (expressed as the *log* of *R^2^*-1), fractal dimension (*D*) and height range (as the *log* of 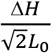) from digital elevation models (DEMs) for 591 reef patches of 4 m^2^ at 21 reef sites encircling Lizard Island on the Great Barrier Reef, Australia (see Methods). Analyses of the structure of these patches reveal that while rugosity, fractal dimension and surface height range are not independent, they have substantial independent variation (*r^2^* for pairwise relationships between surface descriptors ranging between 3% and 30%, Fig. 2a-c). However, when framed together, the three descriptors formed a plane, whereupon the trivially measured surface descriptors, rugosity and height range, captured 98% of the variation in *D* (Fig. 2d). The remaining 2% of the variation occurs because real surfaces do not necessarily behave like fractals (i.e., are self-similar) across a wide range of scales (Extended Data Fig. 5). The observation that the structure of nearly all measured reef patches fell upon a plane delineated by three simple surface descriptors highlights the fundamental geometric constraints of surface habitats. If fractal dimension increases, then either rugosity increases, or height range decreases, or both. All three descriptors are essential for capturing structural complexity because they explain different elements of surface geometry: height range captures patch scale variation, rugosity captures fine scale variation (which sums to surface area), and fractal dimension captures degree of space filling when transitioning from broad to fine scales (Extended Data Fig. 1a).

Different reef locations, with different ecological and environmental histories, occupied different regions on the surface descriptor plane (Fig. 3). For example, one site that was stripped of living coral during back-to-back tropical cyclones^22^ largely occupied the region of the plane where rugosity, fractal dimension and surface height range are all low (Fig. 3a); that is, the patches at this site were closest to a theoretical flat surface. Another site also impacted by the cyclones but left littered with dead coral branches, had similar levels of rugosity and height range, but fractal dimension was relatively high (Fig. 3b). In contrast, a site containing several large colonies of living branching coral had patches with the highest fractal dimension and rugosity, yet the height range of these patches was low (Fig. 3c) reflecting the approximately uniform height of living branching corals in shallow waters where water depth and tidal range constrains colony growth. Meanwhile, a site containing large hemispherical *Porites* corals had patches with large height ranges and high rugosity but lower fractal dimension (Fig. 3d). Three sites contained patches with similar distributions of rugosities (Fig. 3b,d,f), and therefore similar surface areas. However, these sites ranged from smooth reef surfaces with large holes (Fig. 3e) to highly bumpy surfaces with no holes Fig. 3b), demonstrating why rugosity alone does not capture structural complexity and how varying mixtures of structural components dictate habitat complexity^14^.

**Fig. 2.**
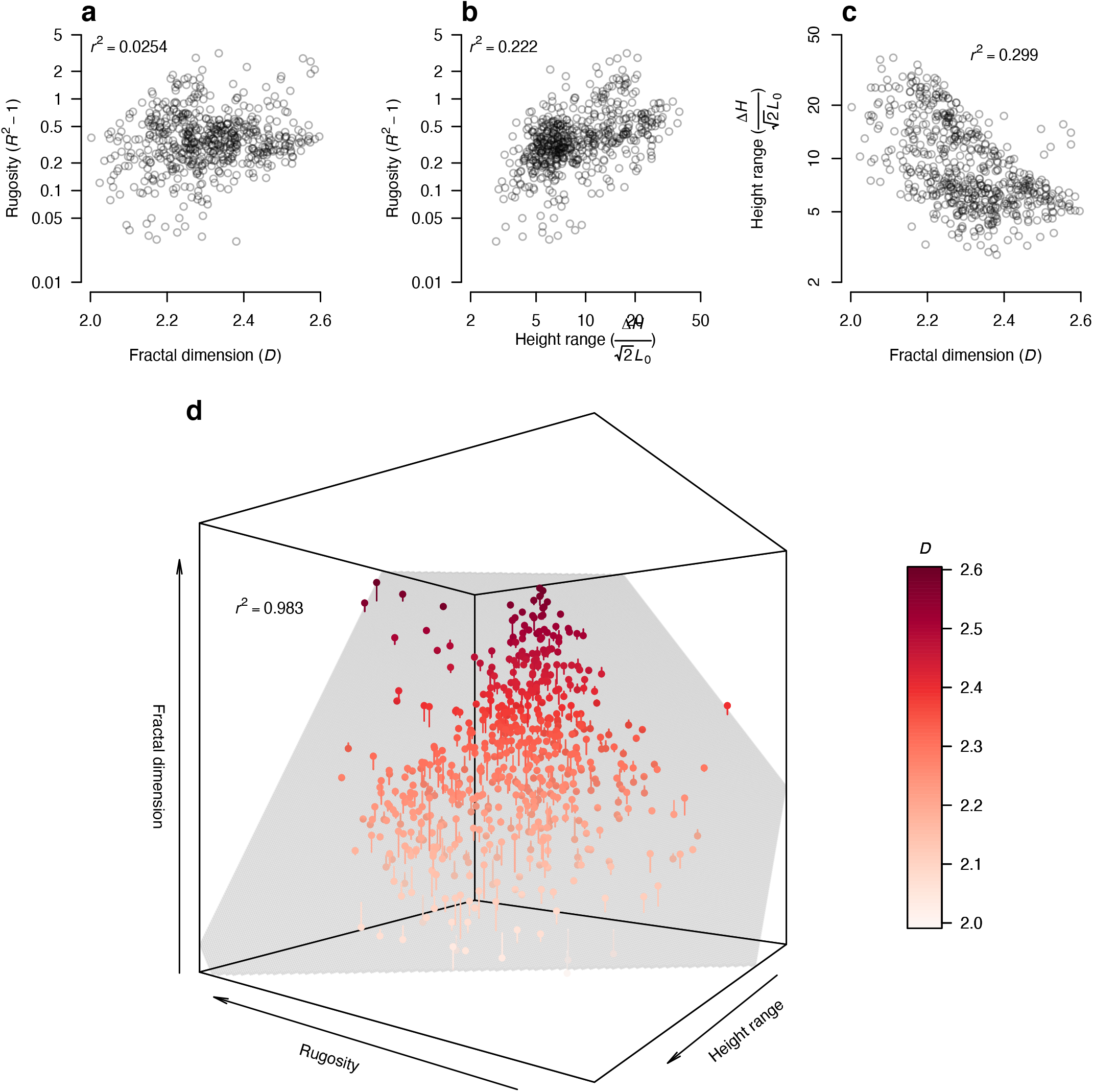
Comparison of the geometric theory with field data. (**a-c**) Pairwise relationships between the descriptors that frame the geometric theory for *n*=595 reef patches: surface rugosity (as *R*^2^-1); fractal dimension *D*; and surface height range (as 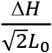). Coefficients of determination (*r^2^*) show the variance explained in the y-axis variable by the x-axis variable. (**d**) When combined the three descriptors explain more than 98% of the variation in fractal dimension *D* despite reef surfaces not being perfectly fractal (see Methods). Field data are points, and the surface descriptor plane is coloured by fractal dimension.

**Fig. 3.**
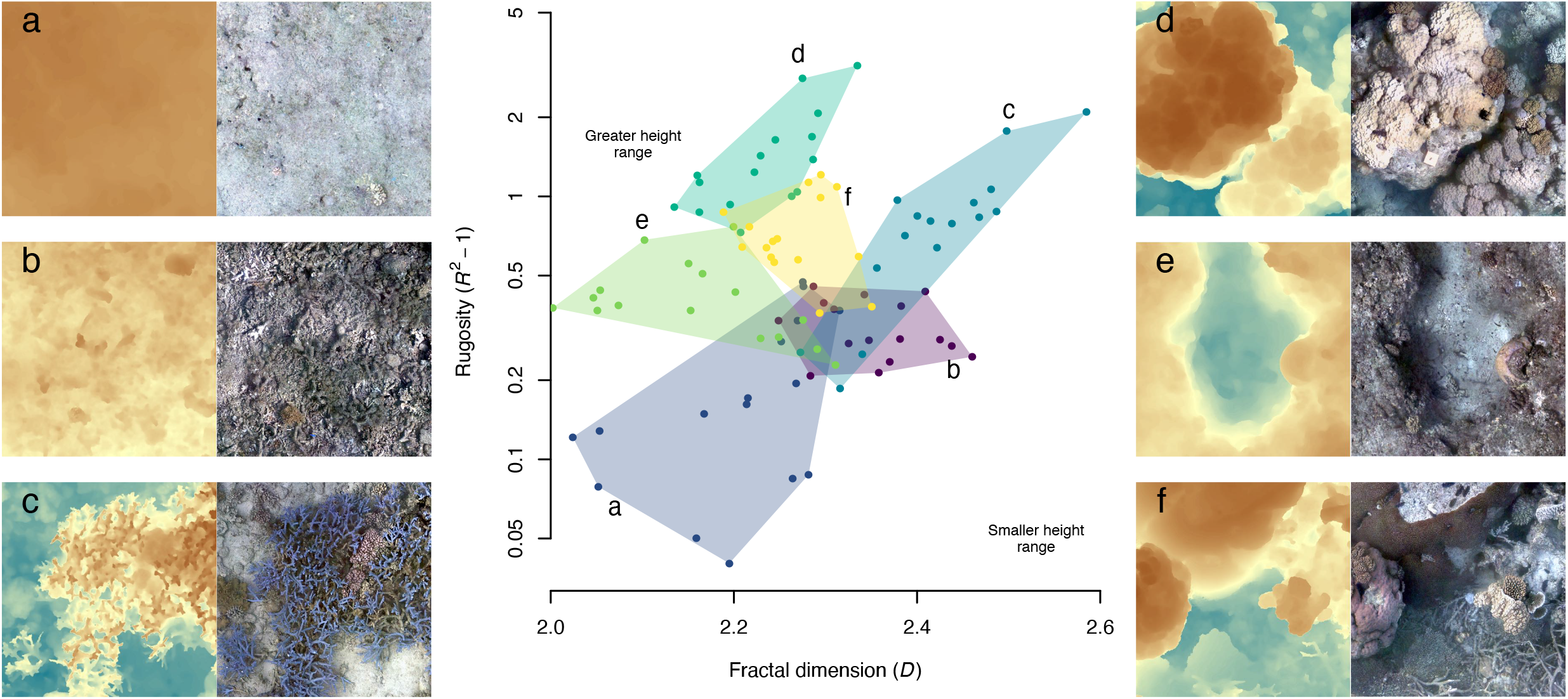
The geometric diversity of coral reef habitats. Reef patches (*n*=16) from a subset of six sites are superimposed onto a two-dimensional representation of the surface descriptor plane (colour used here to delineate sites). (**a**) North Reef; (**b**) Osprey; (**c**) Lagoon-2; (**d**) Resort; (**e**) South Island; and (**f**) Horseshoe. Patch height range is greater in the top left corner and decreases towards the bottom right corner. The corresponding DEMs and orthographic mosaics show selected patches at each site to help visualise geometric differences.

Finally, to connect the geometric variables to biodiversity, we examined how species richness, total abundance and diversity (measured as the probability of interspecific encounter^23^) varied across the surface descriptor plane. Strong ecological feedbacks occur between coral reef habitat structure and coral biodiversity metrics. Coral reef structures are largely created by corals, but their structure is mechanistically affected by environmental conditions such as tidal range, currents, storm impacts and wave exposure. For instance, coral larvae are poor swimmers and are more likely to settle in reef patches with small-scale complexity, because they get entrapped by micro-eddies^24^. At the same time, more intricate coral structures (with higher fractal dimension, *D*) are more likely to be damaged or dislodged during storms that flatten reef patches^25,26^. Species-area theory predicts that species richness and abundances should be highest in patches with the greatest surface area^11^ (i.e., highest rugosity). We predicted that higher fractal dimension would also enhance species richness and abundance, because of niche diversity (i.e., increases in surface area at different scales), and that this effect would be additional to overall surface area. The surface descriptor plane allows estimating the combined effects of not just area, but also niche differentiation associated with fractal dimension and height range^10,15^.

We examined geometric-biodiversity coupling for a large plot, containing 261 of the 4 m^2^ reef patches, in which 9,264 coral colonies of 171 species were recorded (see Methods). Contrary to expectation, we found that all biodiversity metrics considered peaked in reef patches with intermediate surface rugosities (Fig. 4a shows diversity, and Extended Data Table 2 includes species richness and abundance). Indeed, several recent studies have argued that the relationship should be unimodal because, as complexity increases, the amount of area available for individuals to live declines^27,28^. However, biodiversity metrics also tended to increase monotonically in association with patches with higher fractal dimension and smaller height range (Extended Data Fig. 7). The explanatory power of reef geometry on biodiversity metrics was over 50% (Extended Data Table 1)—5 to 45% higher than any surface descriptor alone. Explaining this much variation in biodiversity is striking, given the number of other, nongeometric processes that govern coral biodiversity, including environmental filtering, dispersal and species interactions^29^. Because corals are autogenic ecosystem engineers, reciprocal causality is likely to strengthen and shape geometric-biodiversity coupling. For instance, high rugosity is often generated by large hemispherical corals (e.g., Fig. 3d) that reduce the number of individuals, and hence species, per area^14^. Subsequently, geometric-biodiversity coupling may be weaker for other surface-associated taxa, such as fishes and invertebrates, and should be tested. Nonetheless, our findings have implications for resilience following disturbances and for restoration efforts that aim to maximise biodiversity^30^, specifically identifying the reef structural characteristics that should be maintained (or built) to maximize biodiversity.

**Fig. 4.**
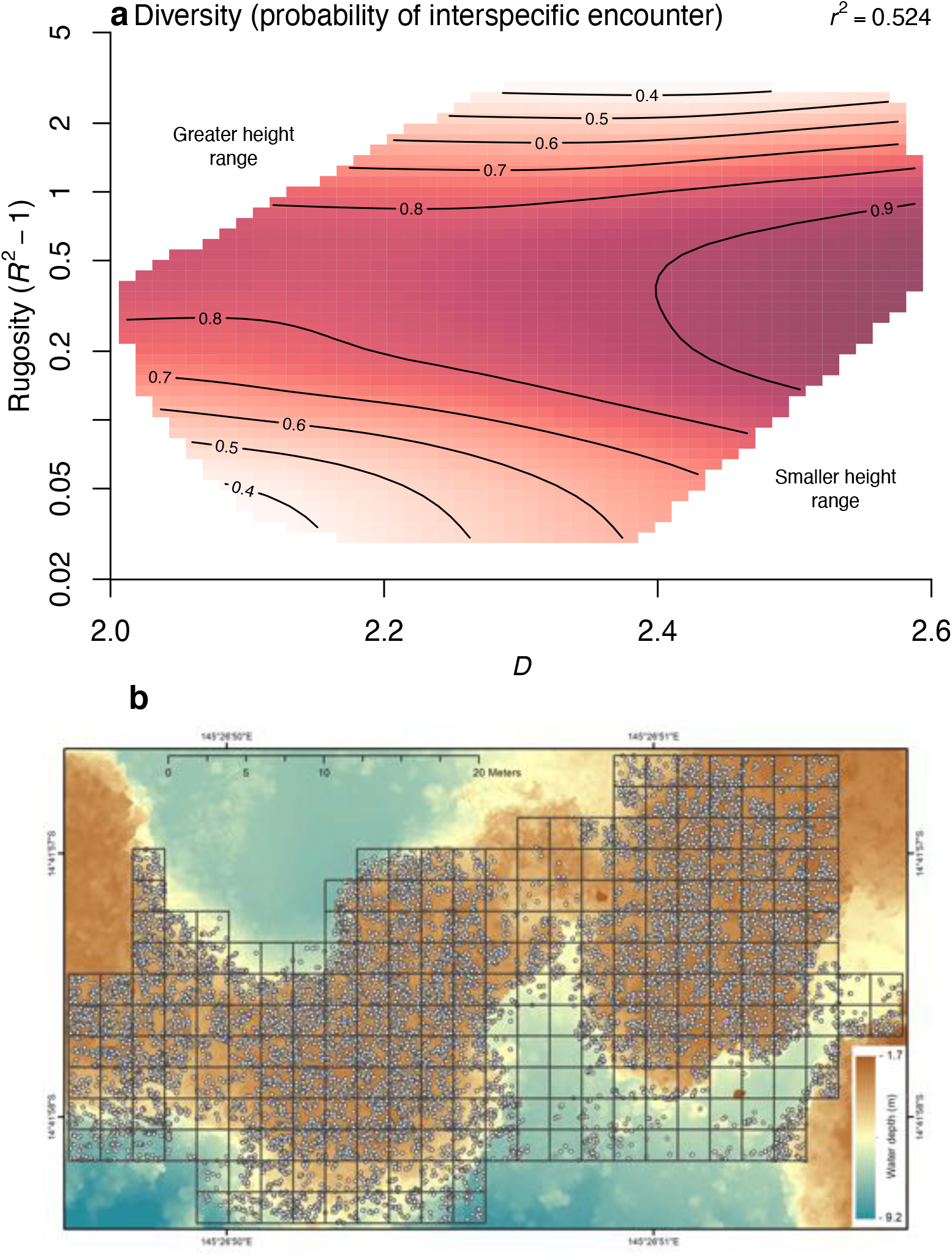
Geometric-biodiversity coupling of coral reef habitats. (**a**) Predicted coral species diversity (represented as probability of interspecific encounter) when plotted upon the surface descriptor plane given by rugosity and fractal dimension (height range is greater in the top left and decreases towards the bottom right, as per Eq. 1). Prediction contours are from the general additive model summarised in Extended Data Table 2. (**b**) A digital elevation model of the large plot with *n*=255 contiguous 2 x 2 m reef patches (black squares) capturing 9,264 coral colony annotations (white points) representing 171 species.

## Discussion

A general, scale-independent geometric basis for surface habitats provides a much-needed way to quantify habitat complexity across ecosystems and spatial scales. Meanwhile, creating three-dimensional habitat surfaces is becoming increasingly accessible and cost effective, for example using Structure from Motion^31,32^, both underwater and on land. The importance of surface complexity as a determinant of habitat condition, biodiversity, and ecosystem function is well recognised^33^, yet different metrics are typically used for different ecosystems, or different taxa within the same ecosystem^10^. The general quantitative approach we propose is applicable across surface habitats in both marine and terrestrial environments, allowing formal comparisons examining whether geometric-biodiversity couplings differ among systems in terms of both pattern and strength. The surface descriptor plane uncovered here clearly defines the fundamental geometric constraints acting to shape surface habitats, and consequently, how changes in surface geometry affect biodiversity. Nonetheless, there remain several unknowns about the surface descriptor plane and its associations with biodiversity metrics that require further exploration. These unknowns range from technical limitations (e.g., how does the theory translate from digital elevation models that exclude overhanging surfaces to 3D surface meshes?) to ecological patterns (e.g., how do different types of structural components, such as different mixtures of branching and hemispherical corals or live and dead elements^14,34^, mediate geometric-biodiversity coupling?).

As powerful ecosystem engineers, humans are modifying the planet through the structures we destroy, both physically and indirectly via environmental change^4^, and those we construct. Indeed, human-modified structures differ significantly in their geometry from nature-built structures^35^. Determining how biodiversity, conservation status and recovery rates relate to habitat complexity measures is paramount in the Anthropocene. The approach we propose here allows for predictions of the biodiversity consequences of these structural changes across land and seascapes.

## Methods

### Geometric theory for surface habitats

The variation method for calculating fractal dimension *D* measures the mean height range of a surface at different scales^36,37^. At the broadest scale, the linear extent *L*, the surface height range is Δ*H* (Extended Data Fig. 1a). At the finest scale, the resolution *L*_0_, the height range (Δ*H_0_*) is the mean of height ranges of all the component areas at that scale. The slope *S* of the resulting log-log relationship (shown in Extended Data Fig. 1a) is:

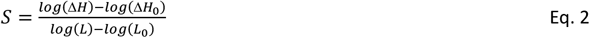

Where fractal dimension is^37^:

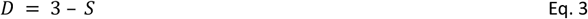

Rearranging Eq. 2 gives:

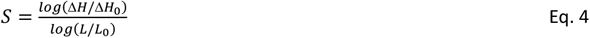

Surface area *A* can be estimated by summing areas *A_0_* at the finest grain *L_0_*. Given the mean height range Δ*H_0_* at *L_0_*, we assume any finer scale detail is not observable, and we calculate *A_0_* from the minimal surface consistent with Δ*H_0_* (Extended Data Fig. 1b) as:

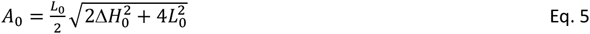

And then multiply by the number of component areas 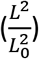 giving:

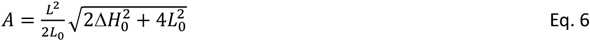

Surface rugosity is^32^:

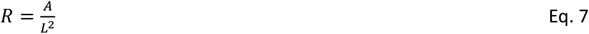

Substituting *A* for Eq. 6 and rearranging gives:

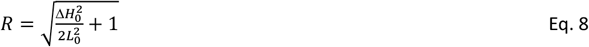

Rearranging for Δ*H_0_* gives:

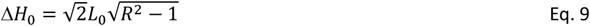

And substituting into Eq. 4 gives:

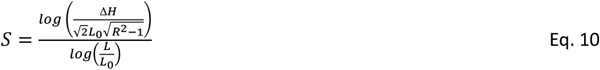

Leading to:

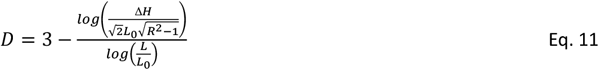

Further rearranging gives a plane as Eq. 1 in the main text. The boundaries equations for the limits of fractal dimension *D* are:

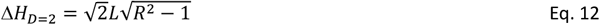

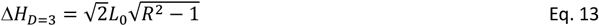

### Coral reef surface field study

Twenty-one reef flat sites were selected approximately 1 km apart and encircling Lizard Island on the Great Barrier Reef, Australia (Extended Data Fig. 2). The spatial arrangement of the sites captured a broad range of habitats that were shaped predominantly by wave exposure generated by prevailing southeast trade winds^22^. Mean water depth across all study sites range between 2 to 3.5 meters. In 2014, at the Trimodal site, we used an Iver2 Autonomous Underwater Vehicle^38^ to collect 45,000 georeferenced overlapping, stereo-pair images of an approximately 30 m by 50 m section of the reef crest (Fig. 4c). In 2016, at all 21 sites, we used the spiral method^39^, which involves swimming a camera rig that unspools from a central point to capture approximately 3000 overlapping, stereo-pair images of approximately 130 m^2^ of reef crest (Extended Data Fig. 3). We used a simultaneous localisation and mapping approach^40^ fusing GPS, stereo imagery and altitude information to provide an initial pose estimate for the cameras. We used Agisoft Metashape software to process the images and produce a 3D dense cloud from which we derived a gridded digital elevation model (DEM) and orthographic mosaic for coral annotation per site. The output resolution of all DEMs was 0.002 m. We used DEMs in order to exclude overhanging surfaces (i.e., only one height for each xy combination), because the degree to which overhangs are captured from plan view photographic surveys is biased by the changing lighting conditions of the environment^41^ (e.g., the sun angle, cloud cover, water turbidity, etc.). On the other hand, plan view surveys were preferred in order to reduce the time costs associated with capturing stereo pairs from multiple view angles over large areas. The use of DEMs will underestimate surface rugosity and fractal dimension; i.e., the reason why *D* tended to range below 2.6. However, given that overhanging structures were rare at our study sites, *R* and *D* measures are likely to exhibit the correct rank order for patches.

Given the lack of coral cover following the 2016 mass bleaching event on the GBR^22^, we used the 2014 Trimodal large plot to quantify geometric-biodiversity relationships (Extended Data Fig. 2). The plot was divided into a contiguous grid of 2 by 2 m reef patches (Fig. 4c, black squares). Patches of the orthographic mosaic were printed on underwater paper and used as reference maps for *in situ* identification of all coral colonies of diameter >5 cm to species by a team of six researchers over four weeks. We focused on the reef crest and flat (shallower areas in Fig. 4c) but also included reef edge and deeper reef. Colonies of unknown or hard to identify species were photographed and identified in consultation with guide books and other observers. Hemispherical *Porites* colonies were identified to genus due to the difficulty differentiating among the few known species without collecting samples for microscopy. Colony annotations were digitized over the orthographic mosaic using QGIS software (e.g., Fig. 4c, white points). Only scleractinian corals were included for analyses. In total, 9,264 coral colonies of 171 species were observed within the 255 reef patches censused. Diversity was calculated as the probability of interspecific encounter (PIE), or 1 – Simpson diversity^23^.

Each of the circular DEMs had a central point, from which an 8 by 8 m square was centred (Extended Data Fig. 3) and divided into 16 contiguous reef patches of 2 by 2 m. DEMs for each 2 by 2 m patch from both the Trimodal large plot (where corals were censused) and the 21 circular plots were cropped to calculate surface rugosity *R*, fractal dimension *D* and the height range of the patch Δ*H*; the latter being the difference between the deepest and shallowest point in a patch. *D* was calculated using the variation method^37^, where each patch was divided into squares with sides lengths (*L*) of 2, 1, 0.5, 0.25, 0.125, 0.0625 and 0.03125 m capturing approximately two orders of magnitude^42^. The resolution *L_0_* for the theory is the smallest scale (i.e., 0.03125 m). The height range within each grid at each scale were calculated, and then averaged for that scale to avoid weighting the many estimates at smaller scales more than the fewer estimates at larger scales when calculating the slope *S*. *S* was calculated for each patch by fitting a linear model to the log of scale (i.e., grid sizes) versus the log of mean height. *D* was then calculated according to Eq. 3. *R* was calculated according to Eq. 8. There are many ways to estimate the surface area of a DEM, so we compared surface rugosity calculated from theory (Eq. 8) with estimates based on surface area calculations using the *surfaceArea* function in the package *sp*^43^. The theory underestimated surface rugosity by approximately 5% (Extended Data Fig. 4), because of the minimal area assumption (Extended Data Fig. 1b), but this disparity was consistent across the range of rugosities.

### Analyses

Surface rugosity (expressed as *R*^2^-1) and standardised height range (expressed as 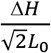) were log-transformed (base 10) as per the plane equation (Eq. 1). Species richness and abundance were sqrt-transformed, and diversity arcsin-transformed, for all analyses to improve model residuals. Coefficients of determination (r square) of pairwise associations of the three geometric variables were estimated by squaring Pearson correlation coefficients. Reef surfaces were not perfectly fractal: mean height ranges at *L* and *L_0_* anchor the theory (Extended Data Fig. 1a), but mean height ranges at scales intermediate to *L* and *L_0_* could shift the overall relationship, albeit subtlety. Therefore, we calculated the r square for the surface descriptor plane based on the deviances of empirically derived *D* from theory derived *D* (Extended Data Fig. 5) (i.e., by dividing the residual sums of squares by the total sums of squares, and then subtracting this value from one).

We quantified geometric-biodiversity relationships for the large plot at the Trimodal site using both generalised additive models (GAMs) and linear models (LMs). We applied the default smoother term to each surface descriptor for the GAMs and second-order polynomials for the LMs, to allow for non-linear relationships among predictor and response variables. We quantified the effect of each geometric variable separately on species richness, total abundance and diversity (PIE), and then all together to assess improvement in explained variation as adjusted r square values (Extended Data Fig. 7; Extended Data Table 1). We included the three reef patches with no living coral, but also confirmed that removing these points had no discernible influence on the geometric-biodiversity relationships. We also ran analyses following the removal of the 5-6 highest rugosity patches that appeared to be largely responsible for producing the hump-shaped rugosity-biodiversity relationships (red curves, Extended Data Fig. 7). Smooth terms for rugosity and height range were significant, with reference degrees of freedom much greater than one for all biodiversity metrics, suggesting significant non-linear effects for these surface descriptors^44^. Fractal dimension showed a linear effect for richness and diversity, and so the smoother term was removed for these analyses. Residuals for all models were approximately normal and were homogeneous when plotted against predictor variables. The linear models with second-order polynomial terms gave the same overall results as GAMs (Extended Data Fig. 7). That is, the polynomial term was significant for the same terms that retained the smoother function in the GAMs. However, the LMs had lower adjusted r square values and so we presented the final results using GAMs (Fig. 4a; Extended Data Table 2).

All analyses, including model selection and diagnostics and figure creation, were conducted in the statistical program language R^45^ and can be downloaded or cloned at GitHub (https://github.com/jmadin/surface_geometry).

## Data and code availability

Source data and code for data preparation, statistical analyses and figures are available at https://github.com/jmadin/surface_geometry.

## Acknowledgments

We thank Dr. Anne Hoggett and Dr. Lyle Vail of the Lizard Island Research Station for their support.

## Author contributions

The study was conceptualized by JSM, DTP, MD and OP. All authors collected the data. JSM and OP developed the theory and JSM ran the analyses. JSM, DTP and OP developed the software pipeline for data and produced the visualizations. The investigation was led by JSM, DTP, MD and OP. JSM and MD led and funded the broader project, with additional field robotics resources provided by OP and SW. JSM wrote the first draft of the paper and all authors reviewed at least one draft.

