## Extended Data for "A geometric basis for surface habitat complexity and biodiversity"

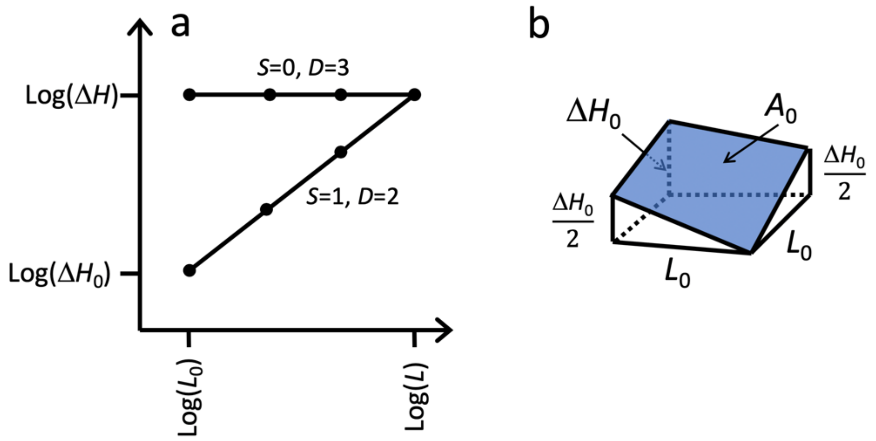


**Extended Data Fig. 1 | Geometric theory for surface habitats.** (**a**) Schematic showing mean height variation as a function of scale. $\Delta H$ is the height range of the habitat surface for the extent $L$. $\Delta H_0$ is the mean height range of the surface at the smallest scale: the resolution $L_0$. The two slopes $S$ represent fractal dimensions $D$ of 2 and 3 according to Eq. 2. For example, high $D$ results when mean height ranges at the scale of the grain are large (i.e., in the vicinity of $\Delta H$), suggesting a highly convoluted surface. Conversely, low $D$ occurs when mean height variations at the scale of the grain are very small, suggesting an approximately flat surface. (**b**) Area $A_0$ at the scale of the grain $L_0$ is calculated as the minimum surface area given the mean height range at this scale.

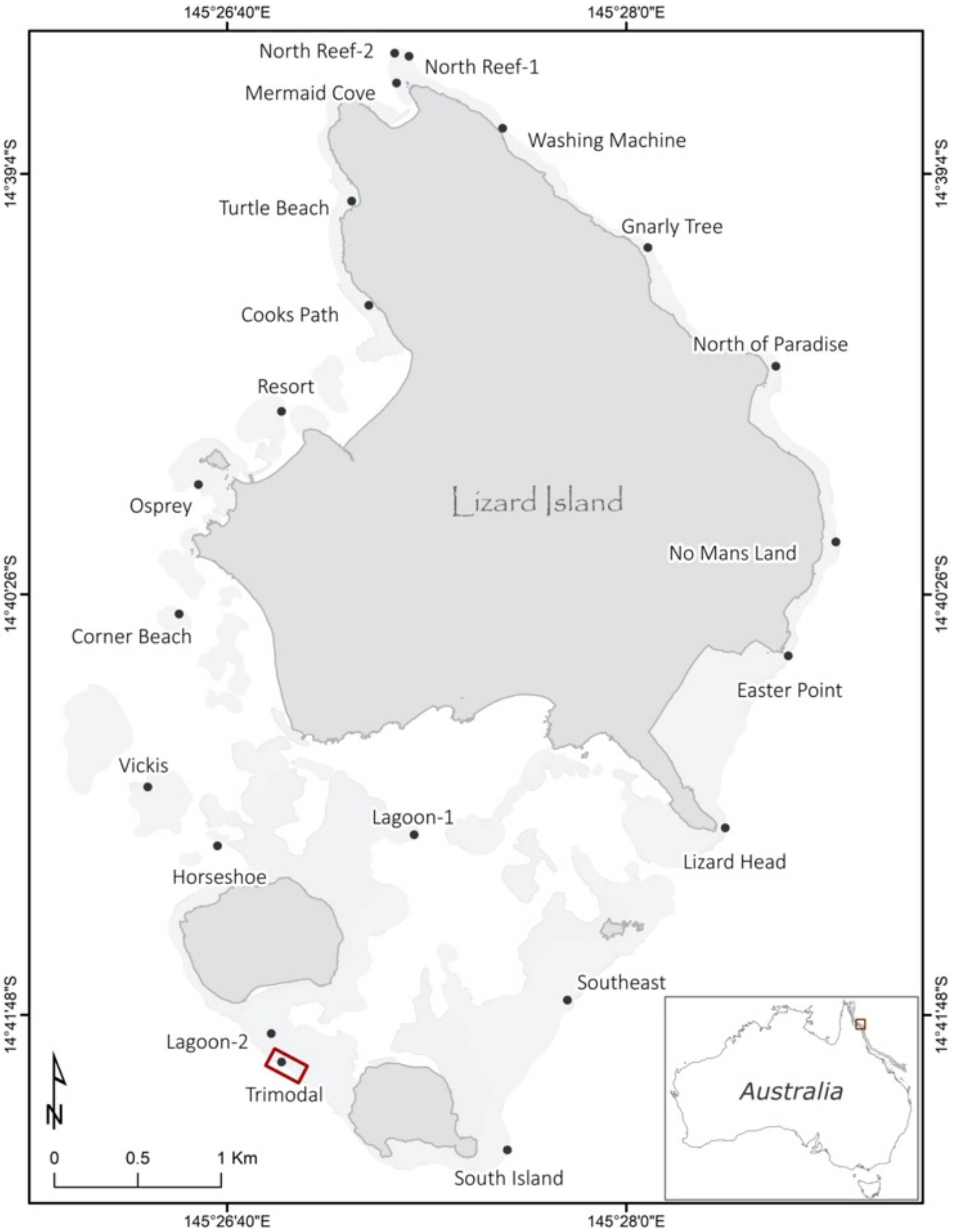


**Extended Data Fig. 2 |** The 21 shallow reef flat locations (black points) and the large Trimodal plot (red rectangle) at Lizard Island, the Great Barrier Reef, Australia.

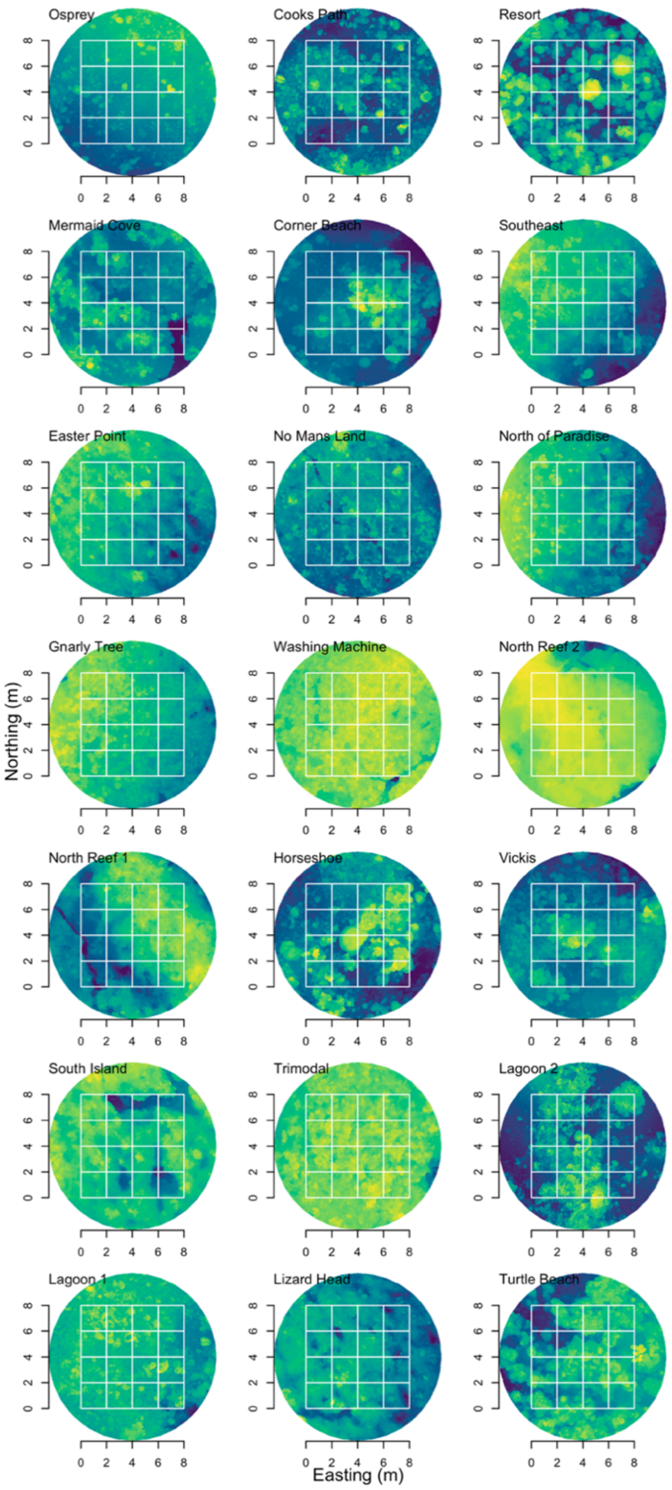


**Extended Data Fig. 3 |** Digital elevation models of the 21 reef sites capturing different habitats encircling Lizard Island (see Extended Data Fig. 2 for map). The models show the 2 m by 2 m reef patches (white squares) and depth gradience from 1.5m (lightest green) to 5m depth (darkest blue).

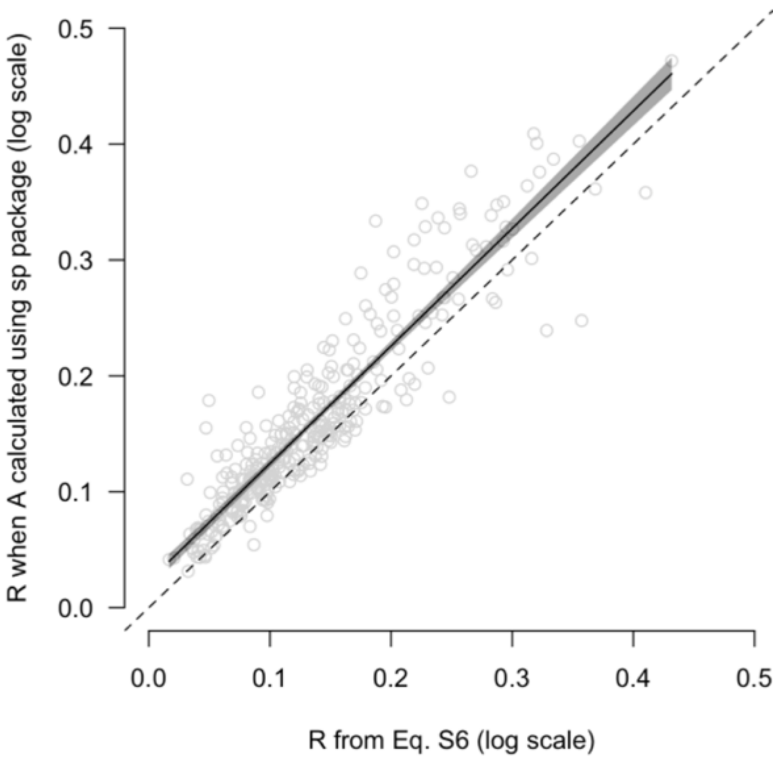


**Extended Data Fig. 4 |** The relationship between *R* when calculated from the geometric theory for habitat surfaces and when calculated using surface area calculated using a spatial statistics function. Dashed line is the unity line. Black line is the fit of a linear model (intercept: 0.023 ± 0.007 95% confidence intervals; slope: 1.014 ± 0.046 95% confidence intervals which encapsulates 1).

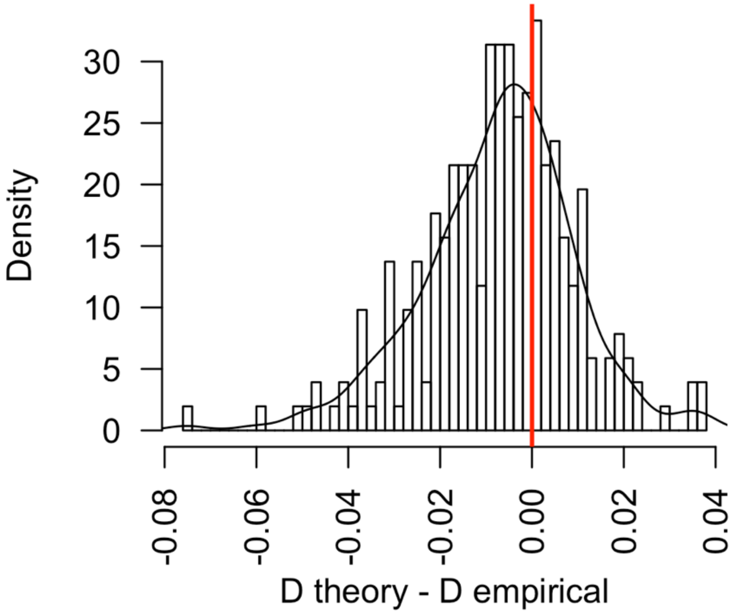


**Extended Data Fig. 5 |** Differences in fractal dimension when calculated empirically versus from the theory that assumes self-similarity across scales.

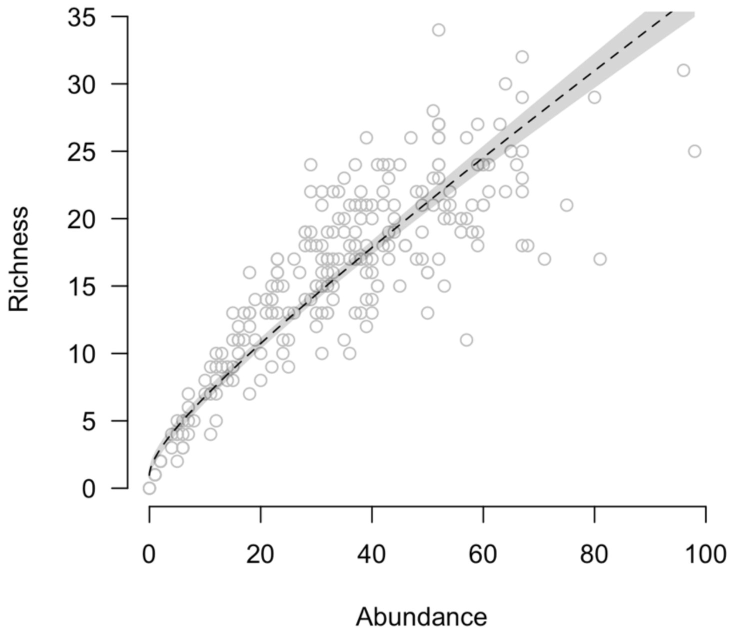


**Extended Data Fig. 6 |** Relationship between abundance and richness within in 2 x 2 m reef patches. Dashed line is linear fit for a model where both axes were sqrt-transformed ($r^2$ = 0.83). Shaded area shows parameter 95% confidence intervals.

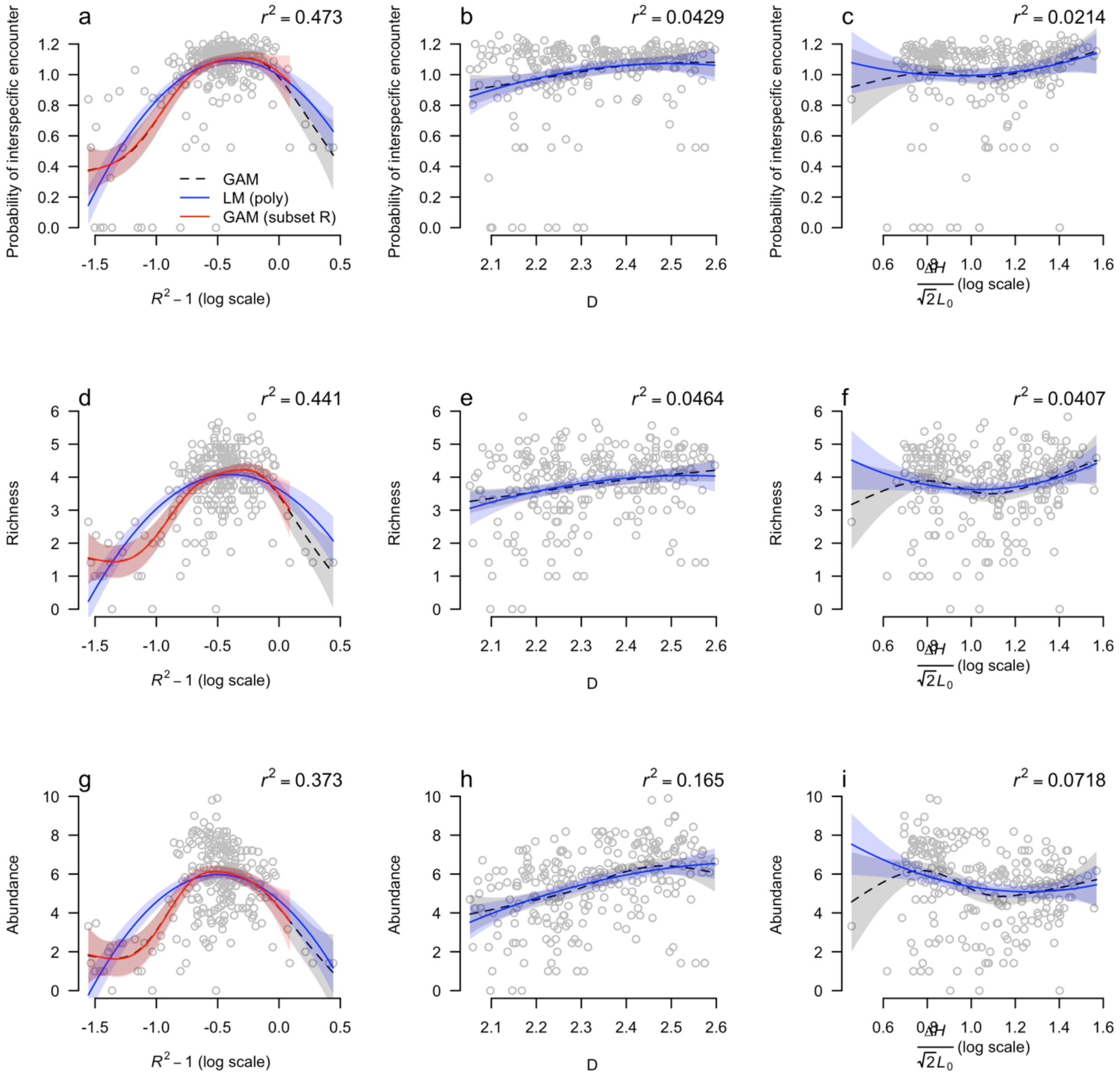


**Extended Data Fig. 7 |** Underlying empirical data and models fits (±95% confidence intervals) for all combinations of the three surface descriptors (x-axes) and species richness, abundance and diversity (PIE) (y-axes). The overall pattern was captured by both statistical approaches: generalised additive models and linear models with second-order polynomials for each surface descriptors. $r^2$ values were consistently better for GAMs and reported here and in Extended Data Table 1.

**Extended Data Table 1 |** Adjusted $r^2$ values for GAM models of (**a**) species richness, (**b**) total abundance and (**c**) diversity (PIE) with each of the three geometric variables and the best combined model after model selection. *N*=255.

| Model | **a** Richness $r^2$ | **b** Abundance $r^2$ | **c** Diversity $r^2$ |
|---|---|---|---|
| $\log_{10}(R^2-1)$ | 0.441 | 0.373 | 0.473 |
| $D$ | 0.046 | 0.165 | 0.043 |
| $\log_{10}(\frac{\Delta H}{\sqrt{2}L_0})$ | 0.041 | 0.072 | 0.021 |
| $\log_{10}(R^2-1) + D + \log_{10}(\frac{\Delta H}{\sqrt{2}L_0})$ | 0.514 | 0.514 | 0.524 |

**Extended Data Table 2 |** Model parameter and smooth term estimates for the best-fit (**a**) species richness, (**b**) abundance and (**c**) diversity (PIE) GAMs. *N*=255.

| **a** Species richness | | | | |
|---|---|---|---|---|
| *Parametric terms* | *Estimate* | *Std. Error* | *t value* | *Pr(>\|t\|)* |
| *D* | 1.624 | 0.021 | 77.23 | << 0.001 |
| *Smooth terms* | *edf* | *Ref.df* | *F* | *p-value* |
| $\log_{10}(R^2-1)$ | 5.496 | 6.651 | 28.016 | << 0.001 |
| $\log_{10}(\frac{\Delta H}{\sqrt{2}L_0})$ | 3.204 | 4.069 | 3.842 | 0.004 |

| **b** Abundance | | | | |
|---|---|---|---|---|
| *Parametric terms* | *Estimate* | *Std. Error* | *t value* | *Pr(>\|t\|)* |
| *D* | 2.346 | 0.037 | 62.92 | << 0.001 |
| *Smooth terms* | *edf* | *Ref.df* | *F* | *p-value* |
| $\log_{10}(R^2-1)$ | 5.695 | 6.860 | 26.875 | << 0.001 |
| $\log_{10}(\frac{\Delta H}{\sqrt{2}L_0})$ | 3.139 | 3.989 | 6.876 | << 0.001 |

| **c** Diversity | | | | |
|---|---|---|---|---|
| *Parametric terms* | *Estimate* | *Std. Error* | *t value* | *Pr(>\|t\|)* |
| *D* | 0.438 | 0.005 | 92.28 | << 0.001 |
| *Smooth terms* | *edf* | *Ref.df* | *F* | *p-value* |
| $\log_{10}(R^2-1)$ | 5.145 | 6.273 | 29.840 | << 0.001 |
| $\log_{10}(\frac{\Delta H}{\sqrt{2}L_0})$ | 2.988 | 3.799 | 3.646 | 0.007 |
